# Pore-forming protein βγ-CAT promptly responses to fasting with capacity to deliver macromolecular nutrients

**DOI:** 10.1101/2022.03.20.485013

**Authors:** Zhi-Hong Shi, Zhong Zhao, Ling-Zhen Liu, Xian-Ling Bian, Yun Zhang

**Affiliations:** Key Laboratory of Animal Models and Human Disease Mechanisms of Chinese Academy of Sciences/Engineering Laboratory of Peptides of Chinese Academy of Sciences, Kunming Institute of Zoology, Chinese Academy of Sciences, Kunming, Yunnan 650201, China; Kunming College of Life Science, University of Chinese Academy of Sciences, Kunming, Yunnan 650204, China; Research Unit of Mitochondria in Brain Diseases, Chinese Academy of Medical Sciences, PKU-Nanjing Institute of Translational Medicine, Nanjing, Jiangsu 210032, China; Institute of Basic Medical Sciences, Chinese Academy of Medical Sciences & Peking Union Medical College, Beijing 100005, China; School of Life Science, Division of Life Sciences and Medicine, University of Science and Technology of China, Hefei, Anhui 230026, China; Center for Excellence in Animal Evolution and Genetics, Chinese Academy of Sciences, Kunming, Yunnan 650201, China

**Keywords:** Pore-forming protein, Fasting, Nutrient, Fatty acid, Albumin

## Abstract

During animal fasting, the nutrient supply and metabolism switch from carbohydrates to a new reliance on the catabolism of energy-dense lipid stores. Assembled under tight regulation, βγ-CAT is a pore-forming protein and trefoil factor complex identified in toad *Bombina maxima*. Here, we determined that this protein complex is a constitutive component in toad blood, that actively responds to the animal fasting. The protein complex was able to promote cellular albumin and albumin-bound fatty acid uptake in a variety of epithelial and endothelial cells, and the effects were attenuated by a macropinocytosis inhibitor. Endothelial cell-derived exosomes containing largely enriched albumin and fatty acids, called nutrisomes, were released in the presence of βγ-CAT. These specific nutrient vesicles were readily taken by starved muscle cells to support their survival. The results uncovered that pore-forming protein βγ-CAT is a fasting responsive element able to drive cell vesicular import and export of macromolecular nutrients.

## Introduction

Animals in nature experience fasting due to the variation of environmental conditions. During fasting, nutrient supply and metabolism switch from carbohydrates to reliance on the catabolism of energy-dense lipid stores (Viscarra & Ortiz, 2013; Secor & Carey, 2016). Endocytosis and exocytosis are fundamental in cells for the exchange of information and materials between the cells and the environments (Klumperman & Raposo, 2014; Wu et al., 2014). These primary cellular processes play key roles in diverse cellular actions, including nutrient acquisition, metabolic adjustment, infection and immunity etc. (Antonescu et al., 2014; Palm & Thompson, 2017; van Niel et al., 2018). Albumin serves as a transport and depot protein for numerous endogenous and exogenous compounds in the circulatory system, including fatty acids (FAs), amino acids and metabolites (van der Vusse, 2009; Fanali et al., 2012; Fung et al., 2018). Particularly, during fasting and in the delivery of lipid nutrients to tissue parenchymal cells, FAs are released from adipose tissues into the bloodstream, where they bind to albumin (van der Vusse, 2009; Fung et al., 2018). Albumin-bound FAs traverse the endothelium and are taken up by the underlying parenchymal cells for energy supply (Young & Zechner, 2013; Kimura et al., 2020; Ko et al., 2020). However, the mechanisms mediating the cellular import and export of albumin and/or albumin bound FAs are incompletely understood (Fung et al., 2018; Ko et al., 2020).

Aerolysins are bacterial β-pore-forming toxins (Dal Peraro & van der Goot, 2016). Proteins with an aerolysin fold, the aerolysin family pore-forming proteins (af-PFPs) or aerolysin-like proteins (ALPs), are found in various animal and plant species (Szczesny et al., 2011; Zhang, 2015; Dang et al., 2017). BmALP1, which is found in the toad *Bombina maxima*, is an af-PFP harboring two βγ-crystallin domains and an aerolysin domain. BmALP1 interacts with the trefoil factor (TFF) BmTFF3 to form a βγ-crystallin-fused af-PFP and TFF complex, named βγ-CAT for its domain composition. This protein complex can form membrane pores (channels) with a functional diameter of approximately 1.5–2.0 nm (Liu et al., 2008; Liu et al., 2021). BmALP3, a paralog of BmALP1, lacks the membrane pore-forming capacity, but it acts as a negative regulator of the assembly of the βγ-CAT complex via redox changes depending on environmental cues (Wang et al., 2020). βγ-CAT acts on sensitive cells that have acidic glycosphingolipids in lipid rafts as receptors in the membranes (Guo et al., 2019) to initiate its cellular actions. The complex acts on endocytic and exocytic systems to form channels in endolysosomes, which promotes cell material import and export through endolysosomal pathways (Zhang et al., 2021). These cellular effects have been shown to be involved in immune defense and tissue repair (Xiang et al., 2014; Li et al., 2017; Gao et al., 2019; Deng et al., 2020; Liu et al., 2021) as well as in toad water maintaining by driving macropinocytosis under osmotic stress (Zhao et al., 2021). Acting as a secretory endolysosome channel (SELC) protein, βγ-CAT and its regulatory network (βγ-CAT-pathway) act as a novel PFP-mediated cell vesicular delivery and transport system, playing multiple roles in the adaptation of toads to diverse environmental conditions (Zhang et al., 2021).

The cellular effects of βγ-CAT raise the possibility that this PFP complex can mediate extracellular nutrient import and export through endolysosomal pathways, which may play a role in trans-cellular nutrient supply as proposed before (Zhang et al., 2021). To test this hypothesis, we studied the potential association between βγ-CAT and the response to fasting in toads, and the ability of this PFP protein to mediate the cellular uptake of albumin and albumin-bound FAs. The results revealed that under fasting conditions, βγ-CAT expression and distribution in toad tissues varied in a pattern consistent with the nutrient supply requirements of the tissues. In various toad cells and mammalian cells, βγ-CAT was found to drive the cellular import of albumin and albumin-bound FAs, which then could be exported by exosome release to support the survival of starved muscle cells. Collectively, our experimental evidence demonstrated the close involvement of βγ-CAT in the toad response to fasting, and its capacity to diver nutrient import and export in vesicular form, which suggested a role in mediating the trans-endothelial cell transport of albumin bound FAs for nutrient supply to tissue.

## Results

### βγ-CAT is a promptly responsive element in toad *B. maxima* fasting

A toad (*B. maxima*) fasting model was used to evaluate the possible involvement of βγ-CAT in toad responses to nutrient deficiency. Toads were fasted for 60 days. The fasting resulted in a ~20% reduction in body weights, which could be partially recovered after 7 days of re-feeding (Fig. 1A). During the fasting, no animal deaths were observed. βγ-CAT was detected in the blood of animals feeding normally as determined by hemolytic activity on human erythrocytes, a sensitive method to detect the presence of this PFP complex (Liu et al., 2008; Wang et al., 2020). The hemolytic activity of toad serum was totally abolished by anti-βγ-CAT and anti-Bm-TFF3 antibodies, indicating that the activity was caused by the βγ-CAT complex (Supplementary Fig. S1A). This result indicated that βγ-CAT is a constitutive component in toad blood. In contrast, 5 nM purified βγ-CAT preparation caused ~65% hemolysis of human erythrocytes (Supplementary Fig. S1B). We then examined the hemolytic activity of toad serum at different time points during fasting to reflect the variation in the amount of the βγ-CAT complex in toad serum. Surprisingly, serum βγ-CAT were found to be rapidly exhausted during the first period of fasting (Fig. 1B). This depletion was indicated by a sharp decrease in toad serum hemolytic activity that was observed on day 6 after fasting, which was approximately 14.5-fold lower than that of the control. However, levels of βγ-CAT toad serum largely increased after day 6 and peaked at day 27, as indicated by the EC_50_ value for the hemolytic activity that was 4.2-fold higher than that of the control. Then, levels of βγ-CAT in the toad serum gradually decreased with extended fasting, and 7 days of re-feeding did not restore βγ-CAT activity (Fig. 1B).

**Fig. 1.**
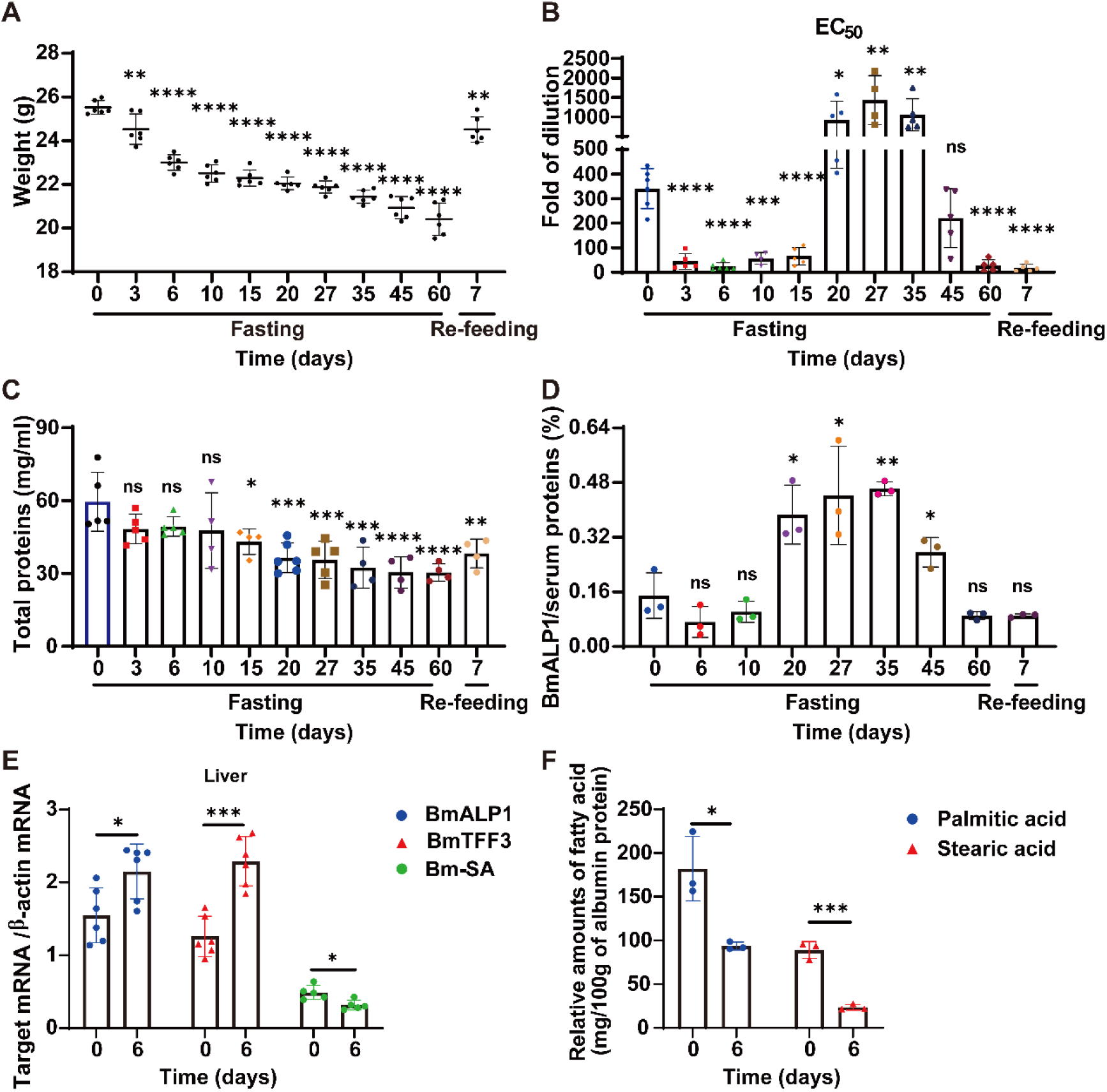
βγ-CAT is a promptly responsive element in toad *B. maxima* fasting. **(A)** Changes in the body weights of toad *B. maxima* during 60 days of fasting and after 7 days re-feeding (n = 6). (**B**) Quantitative comparison of the hemolytic activity of toad *B. maxima* serum at different time points during fasting and re-feeding (n ≥ 4). (**C**) Quantitative comparison of the total protein levels in toad *B. maxima* serum during fasting and re-feeding (n ≥ 4). (**D**) Semi-quantify estimation of the levels of βγ-CAT alpha-subunit BmALP1 in toad *B. maxima* serum. (**E**) The mRNA expression levels of two βγ-CAT subunits and Bm-SA as determined by RT-PCR before and after 6 days of fasting (n ≥ 5). (**F**) Bar graph of the relative FA composition by gas chromatography (GC) of methyl-esterified fatty acid extracts via toad albumin with or without 6 days of fasting. The amount of each lipid class was calculated based on the amount of heptadecanoic acid methyl esters via the peak areas. The data are representative of at least two independent experiments and the data are shown as means ± SD. Statistical analyses were performed with GraphPad Prism 8.0 software: ANOVA for A and C and unpaired t-test for B, D, E, and F. ns: *P* ≥ 0.05, \**P* < 0.05, \*\**P* < 0.01, \*\*\**P* < 0.001, \*\*\*\**P* < 0.0001. In A, B, C, and D, the statistical comparison was made between the data from different time points of the fasting and re-feeding with that of day 0. See also Supplementary Fig. S1.

Next, we examined the changes in the serum total protein levels of the toads during fasting. As expected, extended fasting resulted in a persistent reduction in serum proteins (Fig. 1C). We then determined the serum content of the βγ-CAT alpha-subunit BmALP1 using a semi-quantification method (Supplementary Fig. S1C). In contrast to the persistent reduction in the levels of total serum proteins, the amount of BmALP1 protein showed a variation pattern similar to that observed for the βγ-CAT hemolytic activity, which reached a maximum on day 35, and then progressively decreased (Fig. 1B and 1D). In addition, we analyzed the mRNA levels of βγ-CAT subunits in toad livers on day 6 of fasting. The results showed that although toad albumin (Bm-SA) expression was attenuated because of the fasting, the expression of βγ-CAT subunits in the toad liver was increased (Fig. 1E). The relative contents of FAs (palmitic acid and stearic acid) conjugated to albumin in the toad serum were also assessed. The results showed that albumin-bound FAs were reduced in the toad serum on day 6 of fasting, which was in accordance with the observed decrease in the levels of βγ-CAT on day 6 (Fig. 1F). These results unambiguously uncovered that βγ-CAT is an immediate element in the toad response to fasting.

The organizational and quantitative changes in βγ-CAT complexes in toad muscle and intestinal tissues were analyzed using immunofluorescence to evaluate βγ-CAT distribution during fasting. The large increase in the levels of βγ-CAT in toad muscle detected on day 6 (Fig. 2A and 2B) was consistent with the substantial decrease in βγ-CAT in the toad serum observed on day 6 of fasting (Fig. 1B). These results suggested that the augmented levels of βγ-CAT in toad muscle should be translocated from the serum. The levels of βγ-CAT were found to be largely attenuated in the toad intestine on day 6 of fasting, compared with those of the control (Fig. 2C and 2D). These observations provided further indication of the involvement of βγ-CAT in toad responses to nutrient deficiency.

**Fig. 2.**
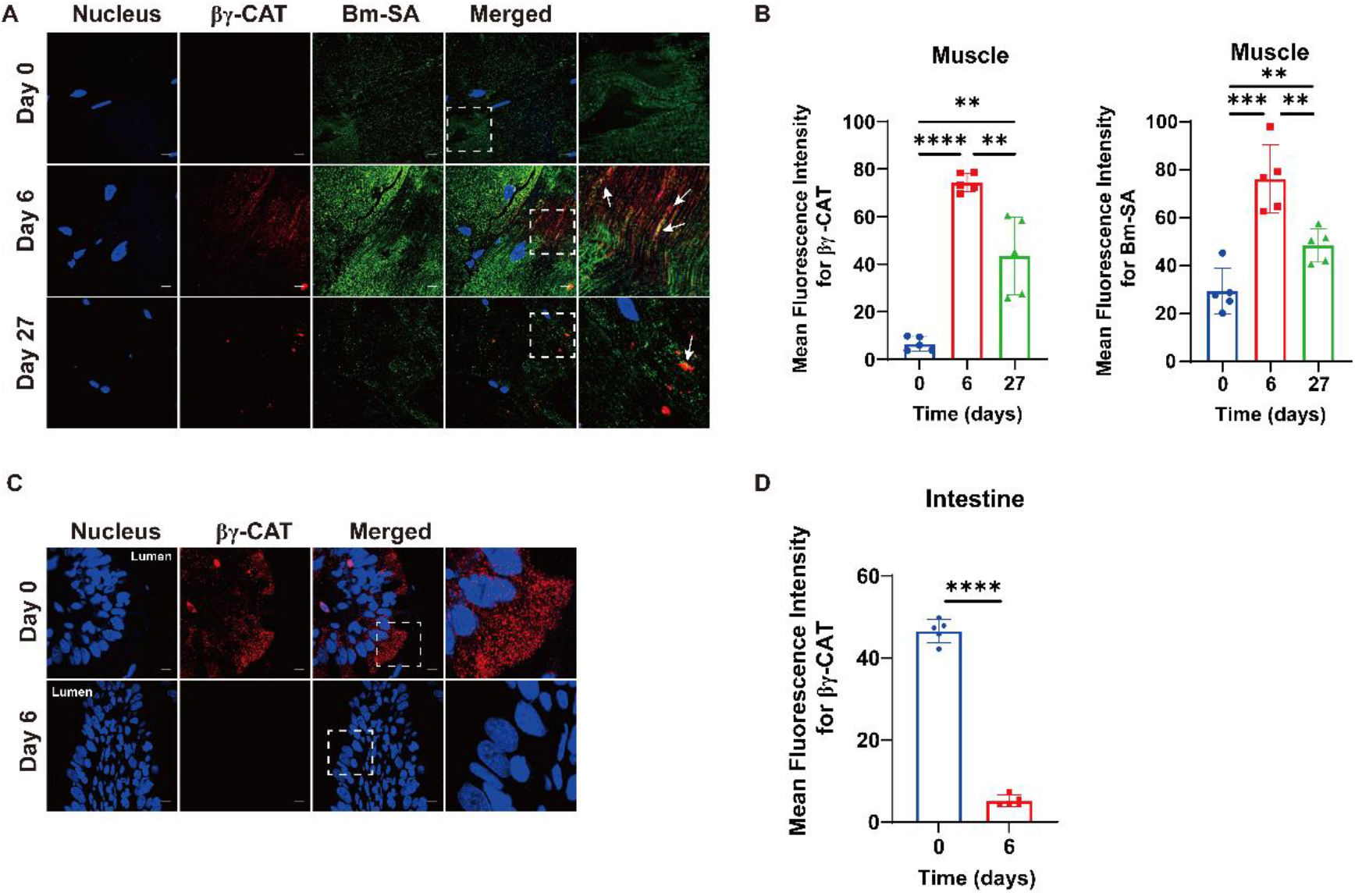
Variation in the distribution of βγ-CAT in toad *B. maxima* tissues under fasting conditions. (**A**) Localization of Bm-SA (green) and βγ-CAT (red) in toad muscle with or without fasting was analyzed by immunofluorescence (IF). Nucleus were stained with DAPI (blue in the image). Scale bar, 10 μm. Up-regulated βγ-CAT and co-localization of Bm-SA and βγ-CAT were observed. Arrows indicate the co-localization of Bm-SA and βγ-CAT. (**B**) Quantitative comparison of the mean fluorescence intensity of βγ-CAT and Bm-SA in (A). (**C**) The localization of βγ-CAT (red) in toad *B. maxima* intestine with or without fasting was analyzed by IF. Nucleus were stained with DAPI (blue in the image). Scale bar, 10 μm. (**D**) Quantitative comparison of the mean fluorescence intensity of βγ-CAT in (C). The data are representative of at least two independent experiments and the data are shown as means ± SD. Statistical analyses were performed using unpaired t-test with GraphPad Prism 8.0 software, \*\**P* < 0.01, \*\*\**P* < 0.001, \*\*\*\**P* < 0.0001.

### βγ-CAT stimulates cellular albumin uptake

We next investigated the possible roles of βγ-CAT in toad responses to fasting. Our previous studies have illustrated that βγ-CAT was able to stimulate the cellular endocytosis of macromolecules via pinocytosis and macropinocytosis, including ovalbumin (antigen) in murine dendritic cells (DCs) and dextran in a variety of different toad cells (Deng et al., 2020; Zhao et al., 2021). Albumin plays important roles in animal responses to fasting, including binding and transporting FAs to tissue parenchymal cells for energy supply (Viscarra & Ortiz, 2013; Secor & Carey, 2016). Thus, we first assayed the ability of βγ-CAT to promote the cellular intake of extracellular albumin in toad skin-epithelial cells and peritoneal cells, as well as in different mammalian cell lines.

First, we purified toad *B. maxima* serum albumin (Bm-SA) through gel filtration on a Sephadex G-100 column. Peak II of the Sephadex G-100 column containing Bm-SA was loaded again on a Q-Sepharose ion exchange column, resulting in the separation of four protein peaks. Bm-SA was found mainly in peak Ⅳ (Supplementary Fig. S2A). We then verified the purity and antibody specificity of purified Bm-SA by Coomassie blue staining and western blotting (Supplementary Fig. S2B). Increased uptake of FITC-Bm-SA and FITC-bovine serum albumin (BSA) by toad skin-epithelial cells and peritoneal cells, respectively, was detected with the addition of 100 nM (for skin cells) or 50 nM (for peritoneal cells) of βγ-CAT as determined by flow cytometry. In contrast, immunodepletion of endogenous βγ-CAT with anti-βγ-CAT antibodies substantially reduced the cellular uptake of the proteins (Fig. 3A and 3B). These results indicated that βγ-CAT promoted the uptake of albumin in toad cells.

**Fig. 3.**
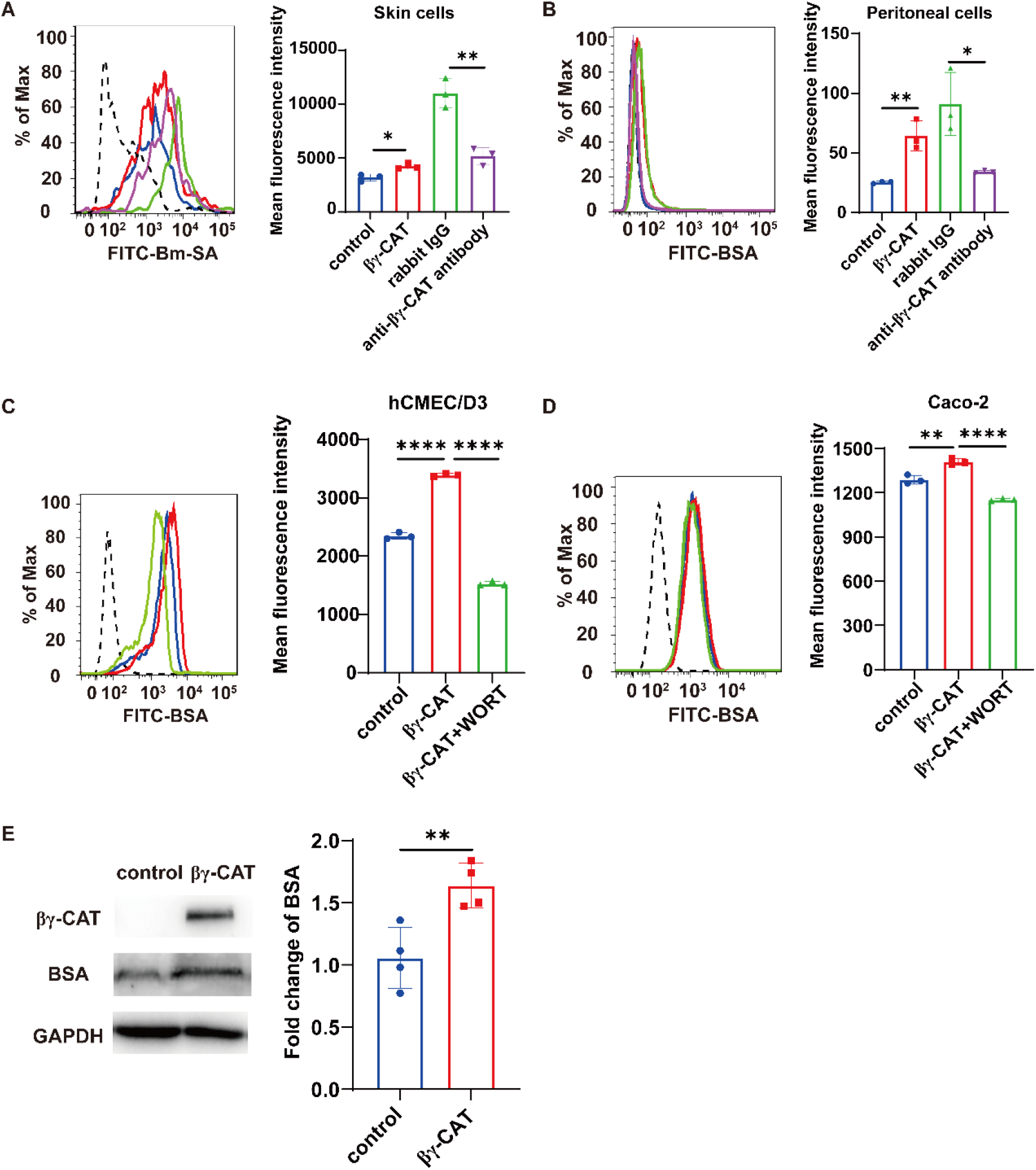
βγ-CAT stimulates cellular albumin uptake. (**A, B**) The addition of purified βγ-CAT increased albumin uptake in toad skin-epithelial cells (A) and peritoneal cells (B). The mean fluorescence intensity of FITC-Bm-SA and FITC-BSA in toad skin-epithelial cells (A) and peritoneal cells (B), respectively, was determined by flow cytometry with or without addition of 100 nM (skin cells) or 50 nM (peritoneal cells) βγ-CAT, for 30 minutes. Immunodepletion of endogenous βγ-CAT decreased albumin uptake. Toad skin-epithelial cells (A) and peritoneal cells (B) were incubated with 100 μg/mL anti-βγ-CAT antibodies for 30 minutes to immunodeplete endogenous βγ-CAT. Then, the mean fluorescence intensity was determined by flow cytometry with 100 μg/mL Bm-SA for toad skin-epithelial cells (A) or 100 μg/mL BSA for peritoneal cells (B) for 30 minutes, respectively. Black dashed line: blank cell group; blue line: control group; red line: βγ-CAT group; green line: rabbit IgG group; and magenta line: anti-βγ-CAT antibody group. (**C, D**) Uptake of BSA by hCMEC/D3 cells (C) and Caco-2 cells (D) was promoted by βγ-CAT and inhibited by the macropinocytosis inhibitor WORT. The hCMEC/D3 and Caco-2 cells were incubated with 100 μg/mL FITC-BSA at 37 °C for 30 minutes with or without 20 nM βγ-CAT and the reactions were followed by the fluorescence detection of FITC. For the inhibition experiment, the cells were first incubated with 20 μM WORT for 1 hour at 37 °C. Black dashed line: blank cell group; blue line: control group; red line: βγ-CAT group; and green line: βγ-CAT + WORT group. (**E**) The intracellular amount of BSA in hCMEC/D3 cells was determined by western blotting after incubation with 1.5 μM BSA with PBS or 20 nM βγ-CAT at 37 °C for 3 hours (left). The western blotting bands were semi-quantified using ImageJ (right). The experiments were performed in triplicate, and representative data are shown as means ± SD. Statistical methods: unpaired t-test for A, B, and E; one-way ANOVA for C and D, with GraphPad Prism 8.0 software, \**P* < 0.05, \*\**P* < 0.01, \*\*\*\**P* < 0.0001. See also Supplementary Fig. S2.

The primary vascular endothelial cells of toad *B. maxima* are extremely difficult to obtain, and therefore we used human brain microvascular endothelial cells (hCMEC/D3) to explore the effects of βγ-CAT on endothelial cells. Human colorectal cancer (Caco-2) cells were also used to investigate the function of βγ-CAT. We first analyzed the cytotoxicity of βγ-CAT toward hCMEC/D3 and Caco-2 cells, and the results showed that 20 nM βγ-CAT or lower dosages did not affect the viability of hCMEC/D3 or Caco-2 cells (Supplementary Fig. S2C). We then assayed the ability of βγ-CAT to stimulate macropinocytosis in hCMEC/D3 and Caco-2 cells. Consistent with our previous findings (Zhao et al., 2021), βγ-CAT greatly increased the internalization of the micropinocytosis marker 70-kDa dextran, which was inhibited by addition of the macropinocytosis inhibitor wortmannin (WORT; a PI3K inhibitor) (Supplementary Fig. S2D). FITC-BSA uptake by hCMEC/D3 and Caco-2 cells was enhanced in the presence of 20 nM βγ-CAT as determined by flow cytometry. However, the FITC-BSA uptake in the presence of βγ-CAT was largely attenuated by WORT (Fig. 3C and 3D). We also validated the promotion of albumin import into hCMEC/D3 cells in the presence of βγ-CAT by western blotting (Fig. 3E). Collectively, these results suggested that βγ-CAT facilitated the cellular uptake of albumin via stimulating macropinocytosis.

### βγ-CAT promotes the cellular uptake of albumin-bound FAs

Under fasting conditions, serum albumin serves as a carrier for the transport of FAs to tissue parenchymal cells for nutrient supply (Spector, 1986; van der Vusse, 2009; Chung et al., 2015; Finicle et al., 2018). The immunofluorescence assay revealed that βγ-CAT promoted the import of intracellular neutral lipids in hCMEC/D3 cells, and co-localization of the protein with intracellular neutral lipids was observed (Fig. 4A). Consistently, βγ-CAT enhanced the cellular uptake of BODIPY 558/568 C12 in Caco-2 cells, and co-localization of the protein with intracellular BODIPY 558/568 C12 was observed (Fig. 4B). Increased endocytosis of albumin-bound BODIPY 558/568 C12 in the presence of βγ-CAT (20 nM) was also detected by flow cytometry in hCMEC/D3 cells (Fig. 4C). Finally, we extracted the total intracellular lipids using fatty acid methyl esters (FAMEs) and verified this conclusion by gas chromatography. The relative content of intracellular stearic acid after 3 h of βγ-CAT treatment was shown to be 1.3 times higher than that of the control (Fig. 4D). Taken together, these results revealed that βγ-CAT facilitated the cellular uptake of albumin-bound FAs.

**Fig. 4.**
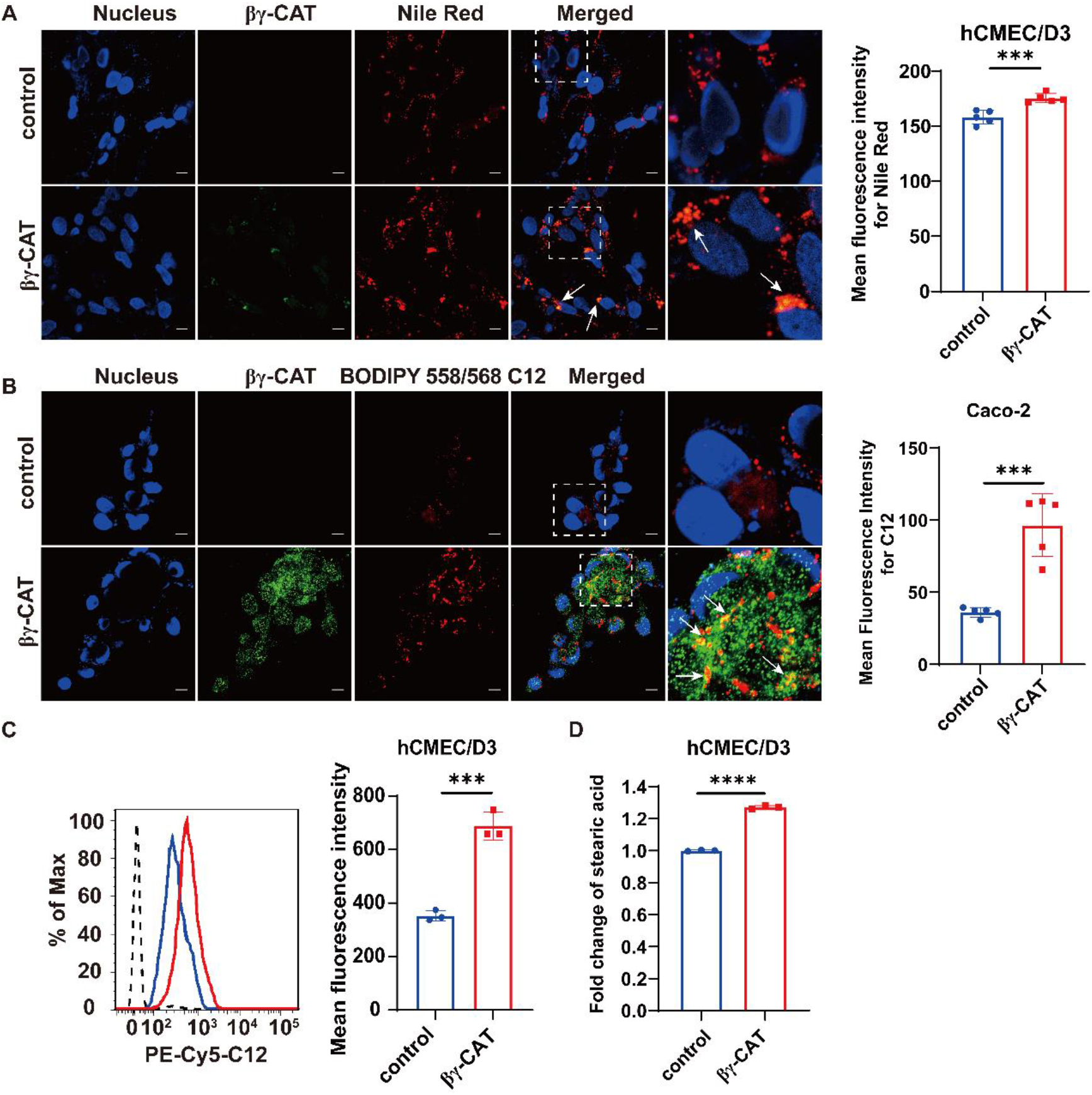
βγ-CAT promotes the cellular uptake of albumin-bound FAs. 5 μM stearic acid or 6.5 μM BODIPY 558/568 C12 was pre-conjugated with 1.5 μM FA-free BSA in 0.01 M PBS at 37 °C for 1 hour. hCMEC/D3 or Caco-2 cells were co-incubated with stearic acid/BSA or BODIPY 558/568 C12/BSA in 0.01 M PBS with or without 20 nM βγ-CAT for 15 minutes at 37 °C. (**A**) The increased import of neutral lipids (stearic acid) (red) in hCMEC/D3 cells in the presence of βγ-CAT (green) and co-localization of the protein with neutral lipids in the cells. Quantitative comparison of the mean fluorescence intensity of intracellular neutral lipids is shown at the right. (**B**) Enhanced uptake of BODIPY 558/568 C12 (red) in Caco-2 cells in the presence of βγ-CAT (green) and co-localization of the protein with BODIPY 558/568 C12 in the cells. The mean fluorescence intensity of intracellular BODIPY 558/568 C12 (right). In (A) and (B), the nucleus were stained with DAPI (blue in the image). Scale bar, 10 μm. Arrows indicate the co-localization of the lipids and βγ-CAT. (**C**) The mean fluorescence intensity of BODIPY 558/568 C12/BSA in hCMEC/D3 cells was determined by flow cytometry with or without 20 nM βγ-CAT treatment for 15 minutes. Black dashed line: blank cell group; blue line: control group; and red line: βγ-CAT group. (**D**) Quantitative comparison of the relative content of intracellular stearic acid in hCMEC/D3 cells co-cultured with stearic acid/BSA and treated with 20 nM βγ-CAT for 3 hours, as determined by fatty acid methyl esters (FAMEs) analysis. The experiments were performed in triplicate, and representative data are shown as means ± SD. Statistical analyses were performed using unpaired t-test with GraphPad Prism 8.0 software, \*\*\**P* < 0.001, \*\*\*\**P* < 0.0001.

### βγ-CAT mediates the export of exosomes containing enriched albumin and FAs

The secretion of exosomes is an important means for the selective export of cellular materials and for intercellular communications (van Niel et al., 2018). βγ-CAT has been shown to stimulate exosome release in murine DCs and a variety of toad *B. maxima* cells (Deng et al., 2020; Zhao et al., 2021). Because βγ-CAT was shown to promote the cellular uptake of albumin and albumin-bound FAs (Figs. 3 and 4) in endothelial hCMEC/D3 cells, we further explored whether βγ-CAT facilitated the export of endocytic vesicles containing albumin and FAs in endothelial hCMEC/D3 cells. The levels of exosome markers CD9, CD63 and flotillin-1 were detected in the collected exosomes (Fig. 5A). Nanoparticle tracking analysis (NTA) revealed that treatment with 20 nM βγ-CAT did not change the average exosome diameter, and no obvious variation in exosome (30–200 nm) concentrations was determined in the presence and absence of βγ-CAT (Fig. 5B). Localization of βγ-CAT was observed in the released exosomes as determined by immunoelectron microscopy (IEM) (Fig. 5C).

**Fig. 5.**
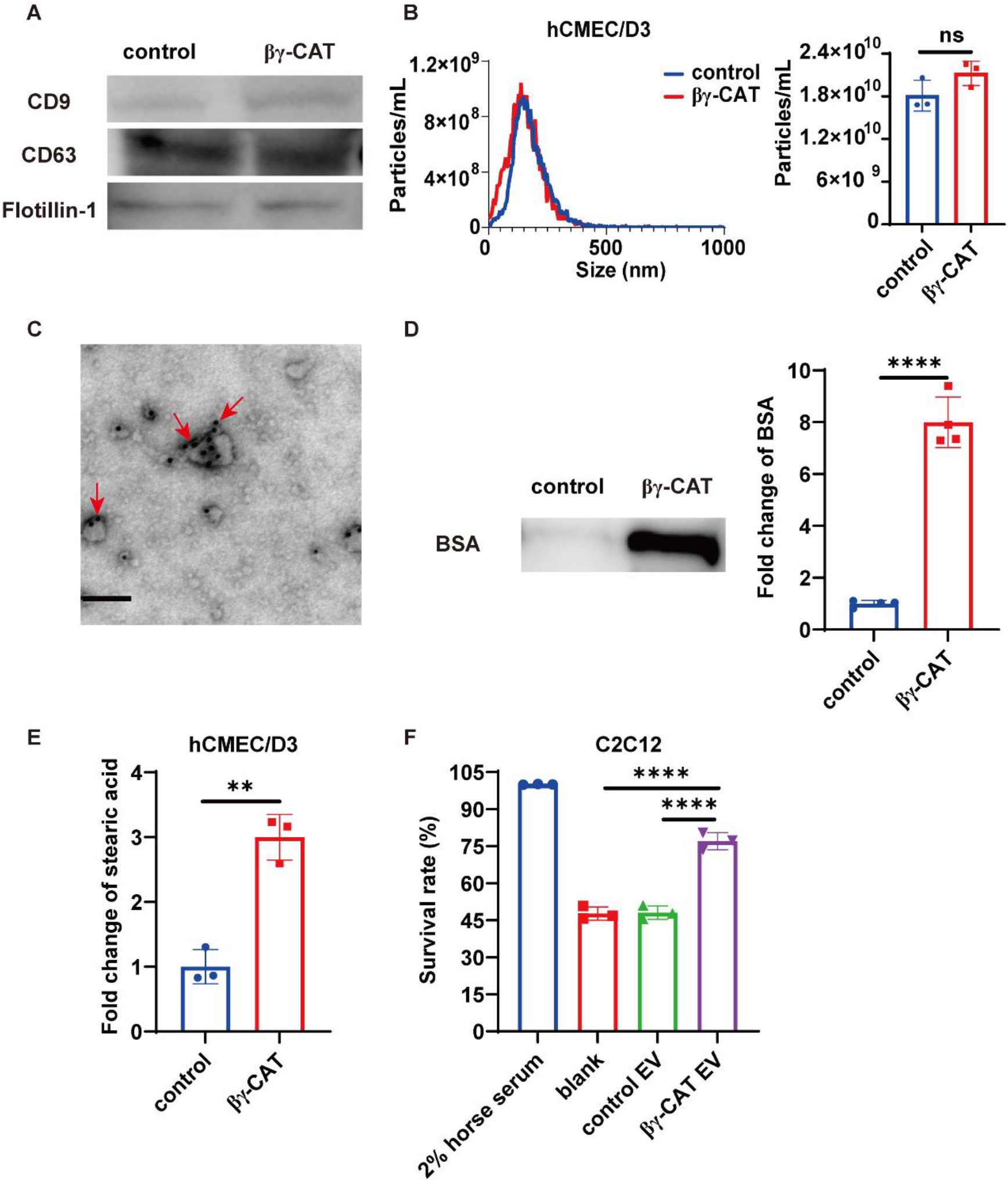
βγ-CAT mediates the export of exosomes containing enriched albumin and FAs. 5 μM stearic acid was pre-conjugated with 1.5 μM FA-free BSA in 0.01 M PBS at 37 °C for 1 hour. hCMEC/D3 cells were co-incubated with stearic acid/BSA in 0.01 M PBS and treated with or without 20 nM βγ-CAT for 3 hours at 37 °C. Then, exosomes produced by the cells were isolated for the following assays. (**A**) The CD9, CD63 and flotillin-1 contents in equal exosome volumes were determined by western blotting. (**B**) Analysis of the concentrations and particle sizes (30–200 nm) of the exosomes from hCMEC/D3 cells by NTA. (**C**) IEM determination of βγ-CAT (as indicated by arrows) in the exosomes from hCMEC/D3 cells. βγ-CAT was labeled with 10-nm colloidal gold particles. Scale bar, 100 nm. (**D**) The albumin content in equal exosome volumes was detected by western blotting. The western blotting bands were semi-quantified using ImageJ (right). (**E**) Quantitative comparison of the relative content of stearic acid in equal exosome volumes as analyzed by FAMEs. (**F**) The cell viability of muscle C2C12 cells determined by MTS. Muscle C2C12 cells were incubated for 12 hours under starvation conditions with or without addition of the exosomes (200 μg/mL) collected from hCMEC/D3 endothelial cells. The experiments were performed in triplicate, and representative data are shown as means ± SD. Statistical methods: unpaired t-test for B, D and E; one-way ANOVA for F, with GraphPad Prism 8.0 software, ns: *P* ≥ 0.05, \*\**P* < 0.01, \*\*\*\**P* < 0.0001.

A large enrichment of albumin and FAs was observed in the exosomes collected from the cells treated with βγ-CAT. The molecular weight of albumin contained in the exosomes from βγ-CAT-treated cells was unchanged, but the amount of the protein was approximately eightfold higher than that of the control, as analyzed by western blotting (Fig. 5D). The lipids were extracted from the exosomes and the relative stearic acid content in the exosomes of βγ-CAT-treated cells was approximately threefold higher than that of the untreated cells as analyzed by the FAME assay (Fig. 5E). These results indicated that βγ-CAT not only was able to stimulate the cellular import of albumin-bound FAs but also mediated the export of albumin and FAs in the form of exosomes. These results were different from our previous findings that when toad *B. maxima* urinary bladder epithelial cells were treated with 50 nM purified βγ-CAT, augmented exosome release was determined but no enrichment of cargo molecules (dextran) was observed (Zhao et al., 2021). This discrepancy clearly reflected that when cells are treated with βγ-CAT, the specific biological outcomes are dependent on the type of cells and the cell surroundings.

To determine whether the exosomes containing albumin and FAs collected from βγ-CAT-treated endothelial cells have biological significance for secondary cells, we investigated the possible role of the exosomes in supporting the survival of muscle cells under starvation conditions. C2C12 cells, which can take up and exploit both serum-derived and cell-derived exosomes to regulate cell growth and differentiation (Vicencio et al., 2015; Mobley et al., 2017), were used in our assays. We found that endothelial hCMEC/D3 cell-derived exosomes collected from βγ-CAT-treated cells were able to increase the survival of muscle C2C12 cells under cell starvation conditions (Fig. 5F). These results elucidated that βγ-CAT mediated the cellular import and export of albumin and albumin-bound FAs, suggesting that βγ-CAT is involved in mediating the transcellular transport of these macromolecular nutrients for toad *B. maxima* tissue requirements, especially under fasting conditions.

## Discussion

Previously, βγ-CAT has been shown to enhance antigen presentation via endolysosome modulation, neutralize endocytic organelle acidification to counteract intracellular pathogens, and cause lysosome destabilization to activate inflammasomes during infection with extracellular pathogens (Xiang et al., 2014; Li et al., 2017; Gao et al., 2019; Deng et al., 2020). Particularly, βγ-CAT promoted the internalization of ovalbumin (an antigen) in a murine DC model via pinocytosis (Deng et al., 2020). βγ-CAT has been shown to facilitates water maintaining in toad by driving macropinocytosis in toad osmoregulatory organs, including the skin and urinary bladder (Zhao et al., 2021). Experimental evidence obtained regarding βγ-CAT and its regulatory network led us to propose the hypothesis of the SELC pathway, which is a novel PFP-driven cell vesicular delivery and transport system (Zhang et al., 2021). The specific βγ-CAT-pathway in toad *B. maxima* is characterized by mediation of the import and export of cellular material (such as water with ions, antigens, and nutrients) through endolysosomal systems. In accordance with our previous hypothesis that βγ-CAT may be involved in nutrient acquisition (Zhang et al., 2021), we found here that βγ-CAT is an immediate element involved in toad responses to fasting (Figs. 1 and 2 and Supplementary Fig. S1). This PFP complex enhanced albumin and albumin-bound FA uptake in various cell lines (Figs. 3 and 4). In addition, βγ-CAT mediated the release of exosomes containing highly enriched levels of albumin and FAs, called nutrisomes. These nutrient enriched vesicles can be taken up by starved muscle cells to increase the survival rate of the cells under starvation conditions (Fig. 5). Dependent on the cell context and surroundings, βγ-CAT is involved in immune defense and water maintaining in toads. In addition, the present study provided evidence that βγ-CAT, a secretory PFP, is active in the fasting state of the toad *B. maxima*, and drived cell macromolecular nutrient import and export. These results suggested that βγ-CAT plays a role in mediating transcellular nutrient transport for tissue energy supply.

mRNAs encoding the two subunits of βγ-CAT have been detected in various toad tissues (Xiang et al., 2014; Li et al., 2017; Zhao et al., 2021). The present study further demonstrated that the assembled βγ-CAT complex was present in toad *B. maxima* blood under normal conditions as indicated by the hemolytic activity on human erythrocytes. Under fasting conditions, blood serum βγ-CAT levels decreased rapidly at the beginning of fasting (Fig. 1B) and the appearance of βγ-CAT in toad muscle was observed (Fig. 2A and 2B). These results suggested that in the beginning of fasting, the rapid exhaustion of toad blood βγ-CAT levels may be caused by the movement of βγ-CAT to toad muscle tissues. The toad weights and serum protein levels continuously decreased during fasting (Fig. 1A and 1C). In contrast, the amount of βγ-CAT alpha-subunit BmALP1in the serum increased in a pattern similar to that of βγ-CAT activity (Fig. 1D). On day 6 of fasting, increased hepatic expression of BmALP1 and BmTFF3 was detected, while decreased albumin expression was observed (Fig. 1E). These results clearly indicated that βγ-CAT is necessary for toad responses to fasting. The observation that βγ-CAT was present in toad intestinal tissue in toads feeding normally, but was absent in fasted toads, further suggested the involvement of βγ-CAT in toad nutrient acquisition and transport (Fig. 2C and 2D).

FAs are basic nutrients for living organisms as energy substrates and as precursors for membrane and signaling lipids (Young & Zechner, 2013; Grasselli et al., 2015; Yen et al., 2015). During fasting, and to deliver lipid nutrients to tissue parenchymal cells, albumin-bound FAs need to traverse the endothelium and be taken up by the underlying parenchymal cells for energy supply (Young & Zechner, 2013; Ko et al., 2020). Several lines of our experimental evidence indicated that βγ-CAT may play a role in facilitating cellular nutrient uptake and transcellular transport during fasting. First, on day 6 of fasting, the levels of both toad serum βγ-CAT and albumin-bound FAs decreased, while albumin in toad muscle increased substantially and was co-localized with βγ-CAT (Figs. 1B, 1D, 1F, 2A, and 2B). This phenomenon suggested that βγ-CAT assists blood albumin in transport of FAs to parenchymal tissues under fasting conditions. Second, our results revealed the ability of βγ-CAT to promote the cellular import of albumin in toad skin-epithelial cells and peritoneal cells as well as in mammalian endothelial hCMEC/D3 cells and epithelial Caco-2 cells (Fig. 3). Third, βγ-CAT enhanced the uptake of FAs in endothelial hCMEC/D3 cells and epithelial Caco-2 cells (Fig. 4). Fourth, highly enriched levels of albumin and FAs were found in βγ-CAT-containing vesicles (nutrisomes) that were exported out of endothelial cells by exosome release, and these nutrisomes could be readily taken up by starved muscle cells (C2C12 cells) to support their survival (Fig. 5). Finally, the variation pattern of βγ-CAT in toad blood and tissues (Figs. 1 and 2) were in accordance with the role of βγ-CAT in cell nutrient delivery for tissue energy supply. From these data, we proposed a model for the action of the secretory PFP, βγ-CAT, in mediating the cellular uptake and transcellular transport of macromolecules for tissue nutrient and energy supply in toad *B. maxima* (Fig. 6).

**Fig. 6.**
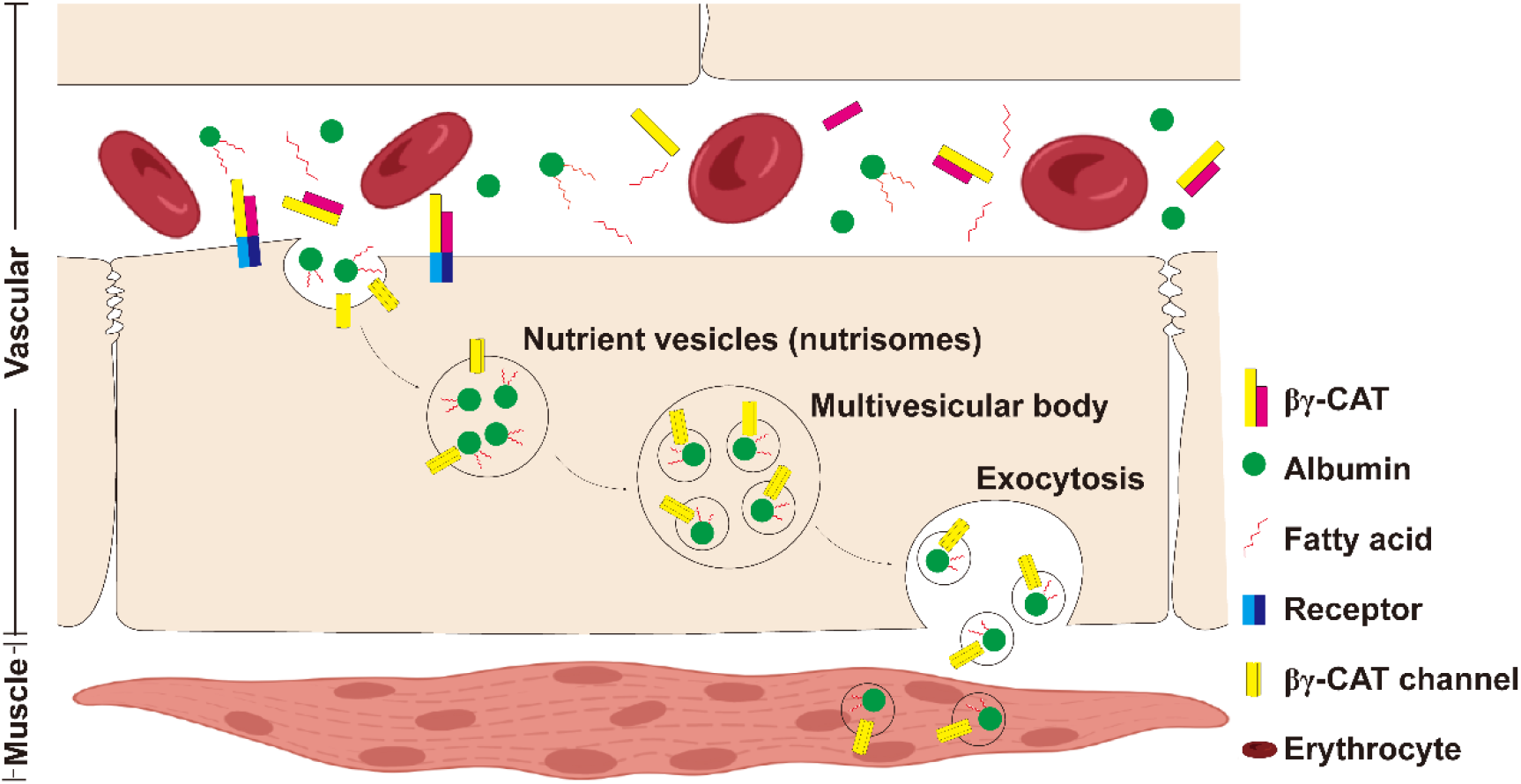
The proposed role of βγ-CAT in mediating cell vesicular delivery of albumin and FAs for tissue nutrient supply. βγ-CAT mediates the cellular uptake of albumin and FAs by promoting endocytosis, probably via macropinocytosis, producing vesicles containing albumin and FAs called nutrisomes. These specific nutrient vesicles (nutrisomes) are exported out of vascular endothelial cells via exosome releasie for nutrient supply to tissue parenchymal cells under fasting. For a detailed description, please see the main text. The presentations of the vascular endothelial and tissue parenchymal cell layers are simplified.

Nutrient transporters, receptor-mediated endocytosis, and macropinocytosis, including pinocytosis, are major types of nutrient uptake pathways (Palm & Thompson, 2017; Palm, 2019). Pinocytosis refers to the cell internalization of extracellular fluid in small endocytic vesicles (< 200 nm), which can supply extracellular proteins to cells (Mayor et al., 2014; Palm, 2019). Previous studies have shown that βγ-CAT promoted pinocytosis in murine DC cells, as determined by Lucifer yellow (LY) uptake assays, which may enhance ovalbumin intake into the cells (Deng et al., 2020). In addition, βγ-CAT has been shown to stimulate macropinocytosis (> 200 nm) in toad skin and urinary bladder epithelial cells, as well as in mammalian MDCK cells under osmotic stress (Zhao et al., 2021). In the present study, the ability of βγ-CAT to promote macropinocytosis was also observed in mammalian endothelial hCMEC/D3 cells and epithelial Caco-2 cells (Supplementary Fig. S2D). The endocytosis of albumin induced by βγ-CAT in these cells was inhibited by the macropinocytosis inhibitor wortmannin (WORT; a PI3K inhibitor) (Supplementary Fig. S2D). These *in vitro* results suggested that βγ-CAT may promote albumin endocytosis through macropinocytosis. However, the possibility that βγ-CAT could facilitate cellular macromolecule import via other forms of endocytic processes, such as receptor-mediated endocytosis or caveolin-dependent endocytosis in distinct cell contexts, cannot be excluded at the present time. Different from classic growth factors that bind to membrane protein receptors to initiate their signaling (Palm, 2019; Swanson & King, 2019), βγ-CAT targets gangliosides and sulfatides as receptors in lipid rafts to promote endocytosis (Guo et al., 2019). The signals downstream from these lipid components and their relationship with cytoskeleton rearrangement require further investigation.

Exosome release driven by βγ-CAT can lead to the export and final transcellular transport of albumin and FAs without disrupting cellular barriers. This mechanism is supported by the observation that enriched levels of albumin and FAs were present in βγ-CAT-containing exosomes (nutrisomes) (Fig. 5D and 5E). This result is consistent with previous studies that indicated βγ-CAT could mediate the export of imported cargo components via exosome release (Deng et al., 2020; Zhao et al., 2021). It has been demonstrated that βγ-CAT can neutralize the acidification of contained endocytic organelles (Xiang et al., 2014; Li et al., 2017; Deng et al., 2020). This cellular process could lead to the translocation of βγ-CAT-containing endocytic organelles into multivesicular bodies (MVBs), which do not fuse with lysosomes for the degradation of contained solutes. This phenomenon was also observed in βγ-CAT-mediated water maintaining in toad urinary bladders (Zhao et al., 2021). βγ-CAT is a PFP protein, and it forms membrane channels with a preference for cations, including Na^+^, K^+^, and Ca^2+^ (Liu et al., 2021; Zhao et al., 2021). The presence of βγ-CAT channels in endolysosomes leads to a rapid flux of Na^+^ (Zhao et al., 2021). This βγ-CAT-induced ion flux might partially explain the acidification modulation observed in the presence of this PFP, which is worthy of further study. It is also possible that some of the albumin-bound FAs imported by βγ-CAT-stimulated endocytosis in endothelial cells might be consumed by the cells themselves as nutrients and for energy supply, which is an interesting topic for future study.

The present study demonstrated that βγ-CAT stimulated toad skin-epithelial cells to uptake albumin (Bm-SA) (Fig. 3A). Amphibian skin has important physiological functions, including being involved in water economy, respiration, metabolite exchange, and immunity etc. (Jørgensen, 1997; Hillyard & Willumsen, 2011; Varga et al., 2018). Toad *B. maxima* albumin is widely distributed around the membranes of epithelial cells and within the stratum spongiosum of the dermis of the skin (Zhang et al., 2005). On one hand, different from albumin circulated in toad blood, there is a haem-b cofactor in albumin accumulated in *B. maxima* skin (Zhang et al., 2005). On the other hand, BmALP3, a paralog of the βγ-CAT alpha-subunit BmALP1, acts as a sensor of oxygen in the toad skin and changes its redox form depending on the oxygen tension, thus, participating in the regulation of βγ-CAT functions (Wang et al., 2020). It is tempting to postulate that βγ-CAT might manipulate the movement of haem-containing albumin, a potential carrier of O_2_, via the import and export of the protein from external environments to internal spaces within multiple epidermal cell layers to facilitate toad skin respiration. In addition, in toad peritoneal cells, βγ-CAT increased the uptake of BSA, an exogenous macromolecule for the toads (Fig. 3B). This result was in accordance with our previous study in which βγ-CAT promoted the internalization of ovalbumin in murine DCs to augment antigen presentation (Deng et al., 2020). Thus, the capacity of βγ-CAT to drive the cellular material delivery in vesicular forms makes this PFP protein complex a versatile multiple functional protein, which has different actions` depending on the cell context and surroundings.

As proposed previously, the SELC protein, βγ-CAT, and its homologous may be viewed as secretory cellular transporters or delivery machines (Zhang et al., 2021), which drive vesicular material delivery to traverse numerous cellular membrane barriers. Accordingly, albumin and albumin-bound FAs are unlikely to be the only macromolecular cargoes taken and transported by βγ-CAT and/or its homologous. Depending on the specific cell type and the cell surroundings, βγ-CAT and/or related homologous may facilitate the import and export of other molecules, including different proteins or non-protein components from the extracellular environment and/or cell membrane systems. The possible existence of specific mechanisms to select particular molecular cargoes in βγ-CAT-stimulated endocytosis is worthy of further investigation in detail. Secretory PFPs have been identified in organisms from all biological kingdoms, and various families of PFPs, including af-PFPs, are widely distributed in plants and animals (Zhang, 2015; Dal Peraro & van der Goot, 2016). Our findings provide a foundation for investigating the roles of PFPs and the mechanisms by which secretory PFPs mediate the cell vesicular delivery of extracellular materials (including nutrients, water containing ions, metabolites, and antigens etc.) through endolysosomal pathways and the related physiological relevance. Particularly, investigating βγ-CAT-like SELC pathways in other vertebrates, especially in mammals, is a fascinating and important future challenge.

In conclusion, the present work elucidated that βγ-CAT, a secretory PFP, is an immediate and active responsive element closely associated with the actions of the toad *B. maxima* facing fasting *in vivo*. *In vitro* results showed that βγ-CAT is able to promote the cellular import and export of albumin and albumin-bound FAs through endolysosomal pathways, which may provide nutrients to secondary cells to support their survival. These results suggested that βγ-CAT may drive the uptake and transcellular vesicular trafficking of macromolecular nutrients to fulfill the biological requirements for energy supply in toad tissues, as proposed (Fig. 6). Our findings provided new insights into the role of secretory PFPs in mediating the transcellular delivery of nutrients for tissue energy supply.

## Acknowledgements

This work was supported by grants from the National Natural Science Foundation of China (grant numbers31572268, U1602225 and 31872226) and the Yunling Scholar Program to Yun Zhang. We would like to thank the Kunming Biological Diversity Regional Center of Instrument, Kunming Institute of Zoology, Chinese Academy of Sciences for electron microscopy observations and we would be grateful to Yingqi Guo for her help of making EM samples. We are grateful to Dr. Fei Li (service center for experimental biotechnology, KIB) for their assistances in GC-MS analysis.

## Conflict of interest

We declare that we have no conflicts of interest.

## Author contributions

Y.Z., Z.H.S and Z.Z. conceived of and conceptualized the study; Z.H.S., Z.Z., L.Z.L., and X.L.B. did the experiments, Z.H.S., Y.Z., Z.Z. analyzed and interpreted the data; Z.H.S and Y.Z. wrote the manuscript; Y.Z., Z.H.S, Z.Z., L.Z.L., and X.L.B. critically revised the manuscript for important intellectual content.

## Materials and Methods

### Animals

Toads (*B. maxima*) were caught in their natural environments. The toads weighing 20-25 g were fed with live *Tenebrio molitor* before the experiments. All procedures and the care and handling of animals were approved by the Ethics Committee of the Kunming Institute of Zoology, the Chinese Academy of Sciences with approval number of **IACUC-OE-2021-05-001**.

### Fasting model of toad *B. maxima*

The toads were placed in a ventilated box with fresh water. The fasting experiment comprised three phases: 3 days of voluntary feeding, 60 days of complete fasting, and 7 days of recovery after fasting. The body weights of the toads were recorded and then the toads were executed. Blood was collected from the heart, which was allowed to stand at 4 °C for 20 minutes, and fresh serum was obtained by centrifugation at 2,000 rpm for 20 minutes.

### Collection and purification of *B. maxima* serum albumin

Purification of toad *B. maxima* serum albumin (Bm-SA) was performed as previously reported (Zhang et al., 2005). The toads were anesthetized with ether, and the blood was collected by a cardiac puncture and was centrifuged at 2,000 rpm for 20 minutes to collect the serum. Briefly, gel filtration using a Sephadex G-100 column resulted in the separation of two protein peaks, and Bm-SA was found in peak II. Peak II of the Sephadex G-100 column was further loaded on a Q-Sepharose ion exchange column and four protein peaks were separated. Bm-SA was found mainly in peak Ⅳ.

### FITC-labeling of Bm-SA

The labeling reactions were performed in accordance with a previously described procedure with some modifications (Giovannoli et al., 2012). Toad serum albumin Bm-SA was dialyzed overnight in a cross-linking solution (NaHCO_3_ 7.56 g, Na_2_CO_3_1.06 g, and NaCl 7.36 g dissolved in 1 L of water; pH 9.0) using a 7,000 Da dialysis membrane (biosharp, Cat BS-QT-020), and 1 mg of FITC (Sigma, Cat F7250) was dissolved in 1 mL of dimethyl sulfoxide (Sigma, Cat D2650). The FITC solution was slowly added to Bm-SA to give Bm-SA: FITC =1 mg:150 μg and the sample was incubated overnight at 4 °C, protected from light. NH_4_Cl (5 M) was added to make the final concentration of 50 mM and the solution was incubated at 4 °C for 2 hours to terminate the reaction. The dialyzed samples were concentrated to 1 mg/mL with PBS and stored at 4 °C in the dark.

### Purification of βγ-CAT

βγ-CAT purification and purity analysis were carried out as described previously (Liu et al., 2008; Xiang et al., 2014).

### Cell culture

As reported previously (Zhao et al., 2021), the primary cells of the toad *B. maxima* were obtained by tissue digestion. Immortalized hCMEC/D3, Caco-2 and C2C12 cell lines were purchased from Kunming Cell Bank, the Chinese Academy of Sciences. hCMEC/D3 cells were cultured in the endothelial cell medium (Sciencell, Cat 1001) containing 5% fetal bovine serum (Sciencell, Cat 0025), 1% penicillin/streptomycin solution (Sciencell, Cat 0503) and 1% endothelial cell growth supplement (Sciencell, Cat 1052). Caco-2 cells and C2C12 cells were cultured in DMEM/F12 (Biological Industries, Cat 01-172-1A) containing 10% fetal bovine serum (Biological Industries, Cat 04-001-1A) and 1% levofloxacin hydrochloride and sodium chloride injection. All cell lines were cultured and grown to confluence in rat-tail collagen type I coated tissue culture flasks (JET Biofil, Cat TCD010100) at 37 °C and 5% CO_2_ in a humid atmosphere.

### Cell viability assay

The MTS assay was performed as described previously (Wu et al., 2020). In brief, hCMEC/D3 and Caco-2 cell lines were seeded in 96-well plates at 1 × 10^4^ cells per well and cultured overnight at 37 °C in 5% CO_2_. Cells were incubated with the MTS reagent (Promega, Cat G3580) in the dark for 1 hour after treatment with βγ-CAT at room temperature for 2 hours. The absorption was detected at 490 nm with an Infinite 200 Pro microplate reader (Tecan, Männedorf, Switzerland).

### Hemolytic activity

Hemolytic activity was determined as described previously (Wang et al., 2020). Serum was collected from toad *B. maxima* at various time points during fasting. Fresh toad serum at different concentrations was mixed with human erythrocytes in equal volumes and incubated for 30 minutes at 37 °C, with saline as a negative control, and 0.5% Triton X-100 as a positive control. After centrifugation at 2,000 rpm for 5 minutes, the absorption at 540 nm was detected with an Infinite 200 Pro microplate reader (Tecan, Männedorf, Switzerland). Calculation of hemolytic activity:

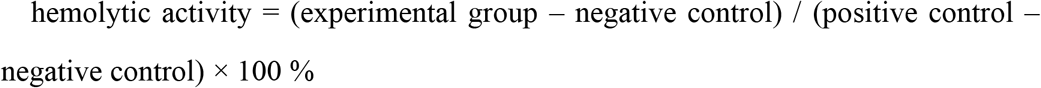

The EC_50_ value was calculated from the logarithmic regression of each concentration–response curve.

For immunodepletion of the endogenous βγ-CAT, the toad serum was incubated with 100 μg/mL rabbit-derived anti-βγ-CAT antibodies or rabbit-derived anti-Bm-TFF3 antibodies for 10 minutes before the above protocol was carried out.

### Protein quantification

Total proteins were measured in the serum of the toads using a BCA protein quantification kit (Thermo Fisher, Cat 23225).

### Quantitative RT–PCR analysis

Quantitative PCR and the primer sequences of βγ-CAT and β-actin were performed as described previously (Xiang et al., 2014). The mRNA levels of BmALP1, BmTFF3, and Bm-SA in the liver were detected by qRT-PCR using a ChamQ Universal SYBR qPCR Master Mix kit (Vazyme, Cat Q711-02). The following primer sequences were used: Bm-SA forward primer: ACGGTATATGCTCACATTGCT and Bm-SA reverse primer: CATGGCAACAGTCCTCGTTC. Target cDNA levels were analyzed by the comparative cycle method and values were normalized against β-actin expression levels.

### Flow cytometry

6.5 μM BODIPY 558/568 C12 (Thermo Fisher, Cat D3835) was pre-conjugated with 1.5 μM FA-free BSA (Meilunbio, Cat MB0094) in 0.01 M PBS at 37 °C for 1 hour (Sun et al., 2020). 2×10^6^ hCMEC/D3 or Caco-2 cells were incubated with 100 μg/mL FITC-BSA (Solarbio, Cat SF063) or 70 kDa FITC-labeled dextran (Sigma, Cat 46945), respectively, in the dark at 37 °C for 30 minutes with and without 20 nM βγ-CAT. The reactions were followed by the fluorescence detection of FITC. 2×10^6^ hCMEC/D3 cells were co-incubated with BODIPY 558/568 C12/BSA solution with or without 20 nM βγ-CAT for 30 minutes in the dark at 37 °C. The reactions were followed by the fluorescence detection of PE-Cy5. For each sample, 1 × 10^4^ single cells were analyzed. 2×10^6^ toad skin-epithelial cells or peritoneal cells were incubated with 100 μg/mL FITC-Bm-SA (skin cells) or 100 μg/mL FITC-BSA (peritoneal cells), respectively, with or without the addition of βγ-CAT (100 nM for skin cells and 50 nM for peritoneal cells) for 30 minutes. For the immunodepletion of endogenous βγ-CAT, the toad cells were incubated with 100 μg/mL rabbit-derived anti-βγ-CAT antibodies for 30 minutes before the above protocol was carried out. For the inhibition experiment, cells were first incubated with 20 μM wortmannin (WORT; a PI3K inhibitor) (Sigma, Cat 681675) for 1 hour at 37 °C. The fluorescence was recorded using an LSR Fortessa cell analyzer (Becton Dickinson, Franklin Lakes, NJ, USA). The data were analyzed by FlowJo 10 and GraphPad Prism 8.0.

### Immunofluorescence

The experiments were performed as previously described with appropriate adjustments (Ktistakis, 2015). Briefly, hCMEC/D3 and Caco-2 cells were grown to 90% on a 24-well glass slide. 5 μM stearic acid (Sigma, Cat S4751) or 6.5 μM BODIPY 558/568 C12 were pre-conjugated with 1.5 μM FA-free BSA in 0.01 M PBS at 37 °C for 1 hour. hCMEC/D3 or Caco-2 cells were then co-incubated with FA/BSA solution with or without 20 nM βγ-CAT for 15 minutes at 37 °C. After washing with PBS three times, the cells were fixed in 4% paraformaldehyde (Servicebio, Cat G1101) for 30 minutes.

The muscle tissues from toad *B. maxima* were collected from non-fasted toads and toads fasted for 6 and 27 days. The intestinal tissues were collected from non-fasted and toads *B. maxima* fasted for 6 days. The blood and contents were washed off with Ringer’s solution, and the tissues were fixed in 4% paraformaldehyde for more than 24 hours.

The fixed tissue sections and cells were treated with 0.5% TritonX-100 for 30 minutes. After blocking for 2 hours in PBS containing 2% BSA at 37 °C, the tissue sections and cells were incubated with mouse-derived anti-βγ-CAT and rabbit-derived anti-albumin primary antibodies (Proteintech, Cat 16475-1-AP) at 4 °C overnight in a dark environment. After washing with PBS three times, the tissue sections and cells were incubated in the dark with fluorescence-labeled secondary antibodies for 2 hours at 37 °C. Coralite488-conjugated goat anti-rabbit IgG (Proteintech, Cat SA00013-2) and coralite594-conjugated goat anti-mouse IgG (Proteintech, Cat SA00013-3) secondary antibodies were used in these experiments. Cellular neutral lipids were stained with 1 μM Nile red (Meilunbio, Cat MB0421) for 15 minutes at 37 °C. The samples were sealed with an anti-fluorescent quench agent containing DAPI (Solarbio, Cat S2110). Images were acquired by a Nikon A1 confocal laser microscope system (Nikon, Tokyo, Japan) or a Zeiss LSM 880 microscope system (Carl Zeiss, Oberkochen, Germany).

### Isolation of exosomes

The isolation of exosomes was optimized based on a previous description (Shao et al., 2018; Zhao et al., 2021). Briefly, hCMEC/D3 cells were grown to 90% on 100-mm cell culture dishes. 5 μM stearic acid was pre-conjugated with 1.5 μM FA-free BSA in 0.01 M PBS at 37 °C for 1 hour. The cells were co-incubated with stearic acid/BSA with or without 20 nM βγ-CAT for 3 hours at 37 °C. The cell supernatant was collected by gradient centrifugation (300 × *g* for 30 minutes; 500 × *g* for 30 minutes; 2,000 × *g* for 60 minutes; and 10,000 × *g* for 60 minutes) at 4 °C. Exosomes were obtained by ultracentrifugation at 100,000 × *g* for 2 hours at 4 °C, then resuspended in PBS, and the exosomes were enriched again using a CP100WX preparative ultracentrifuge (HATACHI, Tokyo, Japan).

### Nanoparticle size analysis (NTA)

The enriched exosomes were diluted in PBS and analyzed for particle size and concentration using a nanoparticle tracking analyzer, Zeta View PMX 110 (Particle Metrix, Meerbusch, Germany), and the data were analyzed with Zeta View 8.04.02 SP2 software.

### Immunoelectron microscopy (IEM)

IEM was performed largely as described previously (Zhao et al., 2021). The enriched exosomes were directly attached to 200-mesh Ni grids (EMCN, Cat BZ10262Na) for 10 minutes before treatment. The samples were blocked with 1% BSA for 5 minutes. After incubation overnight with rabbit-derived anti-βγ-CAT primary antibodies at 4 °C, the samples were incubated with 10-nm colloidal gold-conjugated secondary antibody (Sigma, Cat G7402). The exosomes were stained with 2% uranyl acetate for 3 minutes before observation using a JEM 1400 plus transmission electron microscope at 120 kV.

### Fatty acid methyl esters analysis

A method to prepare fatty acid methyl esters (FAMEs) was used for the analysis of FAs in the serum albumin, cells, and exosomes by gas chromatography. FAMEs were based on the previous report (Maisonneuve et al., 2010; Yang et al., 2016). In brief, 5 μM stearic acid was pre-conjugated with 1.5 μM FA-free BSA in 0.01 M PBS at 37 °C for 1 hour. hCMEC/D3 cells were co-incubated with stearic acid/BSA in 0.01 M PBS with or without 20 nM βγ-CAT for 3 hours at 37 °C. After three washes with PBS, the cells were added to 2 mL of extraction solution (hexane: isopropanol, 1:1). The exosomes in the cell supernatant were enriched following the method of “ **Isolation of exosomes**", and added to 2 mL of extraction solution. Bm-SA from three toads was added to 2 mL of extraction solution. The extracts were ultrasonically crushed and the supernatants were collected by centrifugation at 3,000 rpm for 10 minutes. The FAs in the total lipids were transmethylated using 2 mL of methanol containing 5% H_2_SO_4_ (v/v) with heating at 80 °C for 1.5 hours. In addition, heptadecanoic acid was used as an internal standard at a final concentration of 200 ng/μL for quantification. After cooling, 500 μL of hexane and 2 mL of 1% KCl were added. FAMEs were extracted into the hexane phase by vigorous shaking followed by centrifugation at 2,000 × *g* for 5 minutes. FAMEs were quantified using an Agilent 7890a gas chromatograph/5975c mass selective detector with a 30m DB-5MS capillary column. The MS acquisition parameters included scanning from m/z 50–600 in the electron impact mode for routine analysis. The FAMEs and total lipid contents were calculated by comparing the peak areas.

### Western blotting

Western blotting was performed as described previously (Wang et al., 2020). Gradient amounts of purified βγ-CAT were used for semi-quantitative assessment of the βγ-CAT alpha-subunit BmALP1 in the toad serum. First, a loading volume of 30 μL of purified βγ-CAT containing varying amounts of BmALP1 was prepared. Second, serum from four fasted toads *B. maxima* was mixed and diluted 24-fold to a loading volume of 30 μL. The integrated densities of the βγ-CAT alpha-subunit BmALP1 (38 kDa) bands were analyzed by ImageJ under the same electrophoretic and exposure conditions. The integrated densities of the bands of a series of different purified proteins were graphed as a standard curve to calculate the total amount of BmALP1 protein in the serum of starved toads and the percentage of BmALP1 in the total serum protein.

The levels of βγ-CAT in toad serum, of albumin in cells and of flotillin-1, CD9, and CD63 in exosomes were measured using western blot. hCMEC/D3 cells were incubated with stearic acid/BSA for 3 hours with or without 20 nM βγ-CAT. The cells were collected by centrifugation at 300 × *g* for 30 minutes and the exosomes were obtained according to the “ **Isolation of exosomes**”. The cells and exosomes were washed and lysed for western blotting. Rabbit-derived anti-βγ-CAT, rabbit-derived anti-albumin (Proteintech, Cat 16475-1-AP), mouse-derived anti-GAPDH (Proteintech, Cat 60004-1-Ig), rabbit-derived anti-Flotillin 1 (Proteintech, Cat 15571-1-AP), mouse-derived anti-CD9 (Santa Cruz Biotechnology, Cat sc-13118), and mouse-derived anti-CD63 (Abcam, ab193349) primary antibodies were used in these experiments. HRP-goat anti-mouse IgG (Proteintech, Cat SA00001-1) and HRP-goat anti-rabbit IgG (Proteintech, Cat SA00001-2) secondary antibodies were used in these experiments.

### Incubation of starved C2C12 cells with exosomes from hCMEC/D3 cells

Exosomes from control and βγ-CAT-treated hCMEC/D3 cells WERE incubated with stearic acid/BSA, and isolated and suspended in PBS. C2C12 cells (1 × 10^4^ cells/well) were seeded onto 96-well plates and cultured in the presence of 200 μg/mL exosomes with DMEM medium for 12 h. DMEM medium was used as a negative control and DMEM medium containing 2% horse serum was used as positive control. Finally, the cells were incubated with the MTS reagent (Promega, Cat G3580) in the dark for 1 hour. The absorption was detected at 490 nm with an Infinite 200 Pro microplate reader (Tecan, Männedorf, Switzerland).

### Statistical analysis

Data are expressed as the mean (n ≥ 3) ± standard deviation (SD). The data shown are from single representative experiments, which were repeated independently at least two times for each condition. The standard unpaired t-test was used for comparisons between two groups, and ANOVA was used for comparison of more than two independent groups. Differences with *P* values < 0.05 were considered statistically significant. All statistical analyses were conducted using GraphPad Prism 8.0 software.

---

**Fig. S1.**
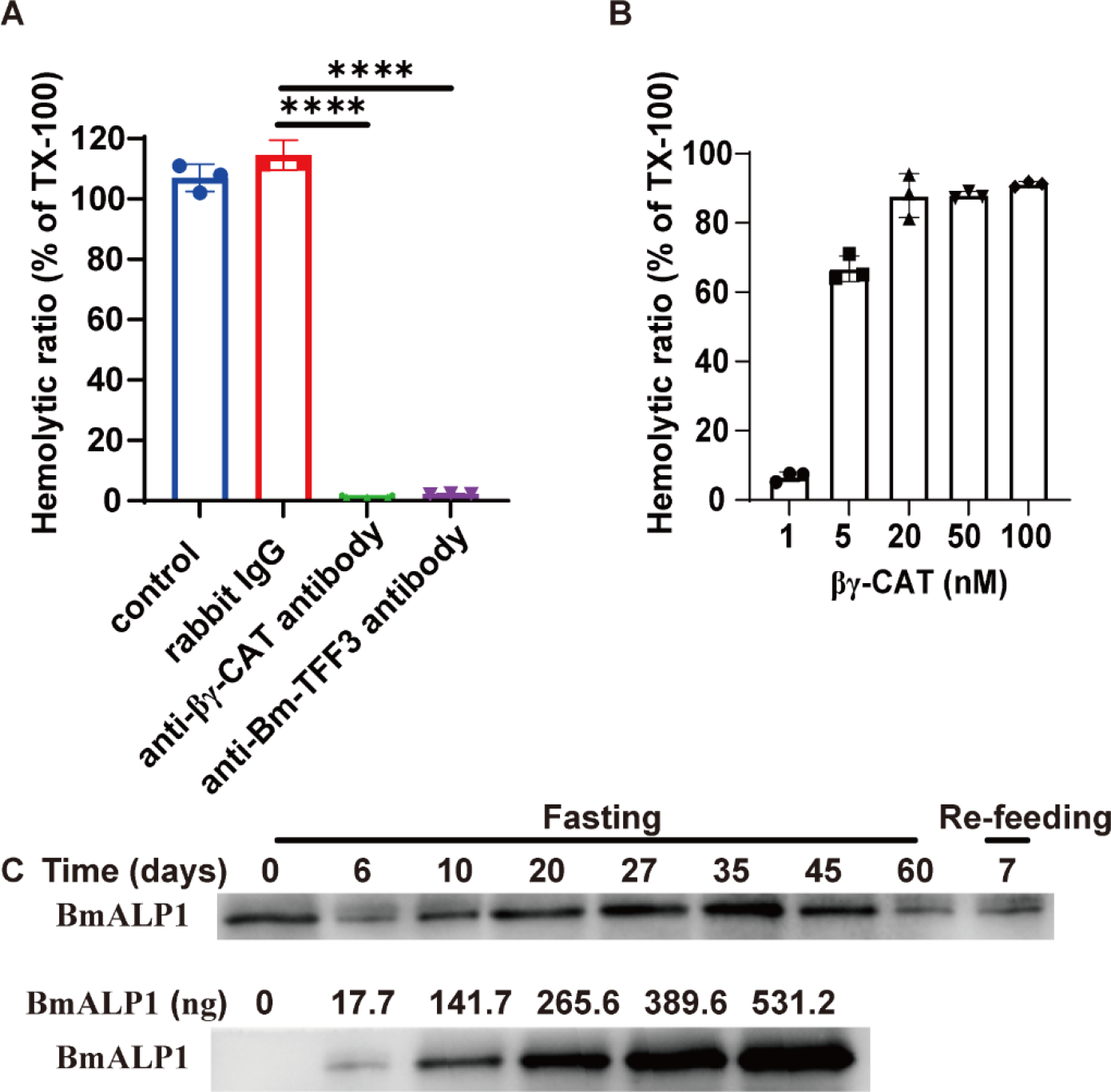
Hemolytic activity of toad *B. maxima* blood βγ-CAT. (**A**) The hemolytic activity of toad serum diluted 100-fold in human erythrocytes was assayed in the absence and presence of 100 μg/mL rabbit-derived anti-βγ-CAT antibodies, rabbit-derived anti-Bm-TFF3 antibodies, and rabbit IgG. Normal saline was used as a control. (**B**) Dosage-dependent hemolytic activity of purified βγ-CAT on human erythrocytes. (**C**) Semi-quantitative assessment of the βγ-CAT alpha-subunit BmALP1 in toad serum diluted 24-fold at various time points during fasting using gradient amounts of purified βγ-CAT. The contents of the βγ-CAT alpha-subunit BmALP1 in toad serum (upper) and purified βγ-CAT (lower) was detected by western blotting. The experiments were performed in triplicate, and representative data are shown as means ± SD. Statistical analysis was performed using one-way ANOVA test with GraphPad Prism 8.0 software, \*\*\*\**P* < 0.0001.

**Fig. S2.**
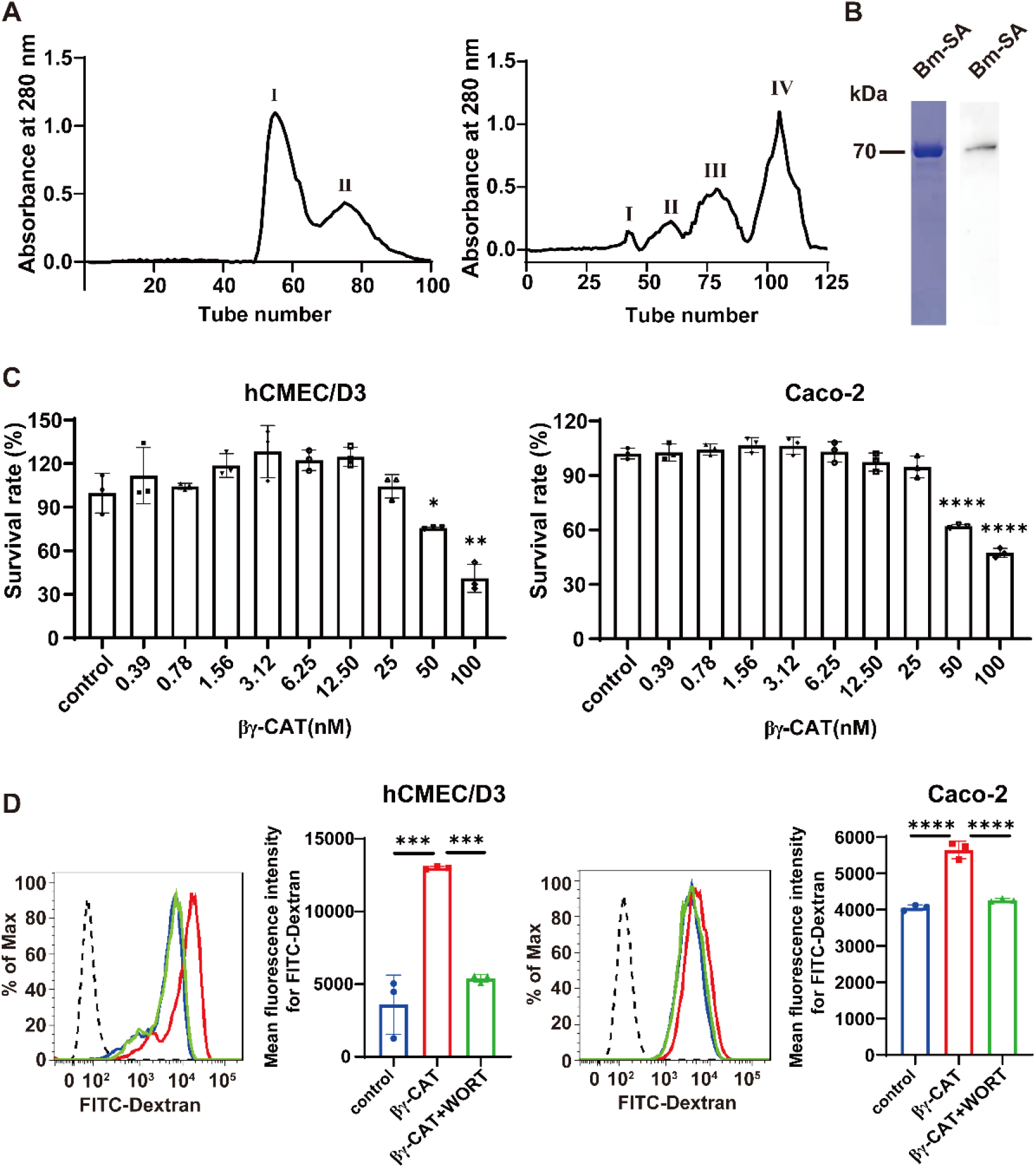
Purification of toad *B. maxima* serum albumin (Bm-SA) and the functional properties of βγ-CAT. (**A**) Purification of Bm-SA. Gel filtration of *B. maxima* serum using a Sephadex G-100 column (left). Peak II of the Sephadex G-100 column was loaded again on a Q-Sepharose ion exchange column. Peak IV was purified Bm-SA. (**B**) The purity of Bm-SA was analyzed by SDS-PAGE with Coomassie blue staining and by western blotting with polyclonal anti-albumin antibodies. (**C**) The cytotoxicity of βγ-CAT on hCMEC/D3 cells and Caco-2 cells. The cells were treated with various dosages of βγ-CAT for 3 hours. Then the cell viability was determined by MTS assay. (**D**) Uptake of 70 kDa FITC-labeled dextran by hCMEC/D3 cells and Caco-2 cells was promoted by βγ-CAT and was inhibited by the macropinocytosis inhibitor WORT. hCMEC/D3 and Caco-2 cells were incubated with 100 μg/mL 70 kDa FITC-labeled dextran at 37 °C for 30 minutes with or without 20 nM βγ-CAT, respectively, and the reactions were followed by the fluorescence detection of FITC. For the inhibition experiment, the cells were first incubated with WORT (20 μM) for 1 hour at 37 °C. Black dashed line: blank cell group; blue line: control group; red line: βγ-CAT group; and green line: βγ-CAT + WORT group. The experiments were performed in triplicate, and representative data are shown as means ± SD. Statistical analysis was performed using unpaired t-test in C and using one-way ANOVA test in D with GraphPad Prism 8.0 software, \**P* < 0.05, \*\**P* < 0.01, \*\*\**P* < 0.001, \*\*\*\**P* < 0.0001.

## Notes

### Competing Interest Statement

The authors have declared no competing interest.

